# High diversity, turnover, and structural constraints characterize TCR α and β repertoire selection

**DOI:** 10.1101/428623

**Authors:** Larisa Kamga, Anna Gil, Inyoung Song, Ramakanth Chirravuri, Nuray Aslan, Dario Ghersi, Lawrence J. Stern, Liisa K. Selin, Katherine Luzuriaga

## Abstract

Recognition modes of individual T-cell receptors (TCR) are well studied, but how TCR repertoires are selected during acute through persistent human virus infections is less clear. Here, we show that persistent EBV-specific clonotypes account for only 9% of unique clonotypes but are highly expanded in acute infectious mononucleosis, and have distinct antigen-specific public features that drive selection into convalescence. The other 91% of highly diverse unique clonotypes disappear and are replaced in convalescence by equally diverse “de-novo” clonotypes. These broad fluctuating repertoires lend plasticity to antigen recognition and potentially protect against T-cell clonal loss and viral escape.

## Introduction

Over 95% of the world’s population is persistently infected with Epstein Barr virus (EBV) by the fourth decade of life. Primary infection commonly occurs in young childhood and is asymptomatic or only mildly symptomatic; primary infection in late childhood or early adulthood often results in acute infectious mononucleosis (AIM)^1^, which is associated with an increased risk of subsequent multiple sclerosis (MS)^2^ or Hodgkin’s lymphoma^3^. EBV infection is also associated with Burkitt lymphoma, nasopharyngeal cancer, hairy leukoplakia in individuals with AIDS, and lymphoproliferative malignancies in transplant patients^3,4^. EBV-associated post-transplant lymphoproliferative disorders can be prevented or treated by adoptive transfer of EBV-specific CD8 T-cells^4–6^. Altogether, these data indicate that EBV-specific CD8 T-cells are important for viral control^7^. Defective CD8 T-cell control of EBV reactivation may also result in the expansion of EBV-infected, autoreactive B cells in multiple sclerosis (MS)^8^; improvement of MS has followed infusion of autologous EBV-specific CD8 T-cells^2^. Improved T-cell therapies would benefit from insights derived from approaches that integrate computational biology and structural modeling to predict optimum TCR features and identify TCR antigen-specificity groups^9–14^. These methods need to be based on an accurate and in depth understanding of antigen-specific TCR repertoire structure and organization from studies like the one presented here. Ultimately, this could lead to better understanding of how EBV-specific CD8 T-cells control EBV replication and facilitate the development of a vaccine to prevent or immunotherapies to modify EBV infection^5,6,15^.

One of the hallmarks of CD8 T-cells is epitope-specificity, conferred by the interaction of the TCR with virus-derived peptides bound to host MHC (pMHC)^16–19^. The TCR is a membrane-bound, heterodimeric protein composed of α and β chains. Each chain arises from a random rearrangement of variable (V), diversity (D), joining (J) and constant (C) gene segments^20^. This recombination process results in a diverse pool of unique TCRα and β clonotypes. Additions or deletions of N-nucleotides at the V(D)J junctions, specifically at the complementarity-determining region 3 (CDR3) and pairing of different TCRα and β segments further enhance the diversity of the TCR repertoire, estimated to range from 10^15^–10^20^ unique potential TCRαβ clonotypes^21,22^. This diversity enables CD8 T-cell responses to a myriad of pathogens.

The CD8 TCR repertoire is an important determinant of CD8 T-cell-mediated antiviral efficacy or immune-mediated pathology^14,21,23–26^. Defining the relationships between early and memory CD8 TCR repertoires is important to understanding structural features of the TCR repertoire that govern the selection and persistence of CD8 T-cells into memory. Deep sequencing techniques, combined with structural analyses, provide a high throughput and unbiased approach to understanding antigen-specific TCRαβ repertoires. We^27^ and others^28–31^ have recently reported that TCRαβ repertoires of CD8 T-cell responses to common viruses (influenza, cytomegalovirus, hepatitis C virus) are highly diverse and individualized (*i.e*. “private”) but “public” clonotypes (defined as the same V, J, or CDR3 aa sequences in many individuals) are favored for expansion, likely due to selection for optimal structural interactions.

To thoroughly evaluate molecular features of TCR that are important for driving repertoire selection over time following EBV infection, we used direct *ex vivo* deep sequencing of TCR Vα and Vβ regions, along with paired TCR Vα/β single-cell sequencing of CD8 T-cells specific to two immunodominant epitopes, GLC and YVL, isolated from peripheral blood during primary EBV infection (AIM) and 6 months later in convalescence (CONV). Each TCR repertoire had a high degree of diversity. However, clonotypes that persisted from AIM to CONV, had distinct public features driving their selection, which was dependent on the specific antigen; while these persistent clonotypes accounted for only 9% of the unique clonotypes, they predominated in both the acute and convalescent phases of infection. The corollary of this finding was that 91% of the unique clonotypes expanded in acute infection were replaced in 6 months by an equally diverse set of de novo clonotypes.

## Results

### Patient characteristics

Four HLA-A*02:01+ individuals presenting with symptoms of AIM and laboratory studies consistent with primary infection were studied (**Table S1**) at initial clinical presentation (AIM) and 6–8 months later (Convalescence; CONV)). Direct tetramer staining of peripheral blood revealed that 2.7%±0.7 (mean+SEM) and 1.3%±0.3 of CD8 T-cells were YVL- and GLC-specific, and declined to 0.3%±0.7 and 0.3%±0.1, respectively, in CONV. Mean blood EBV load was 3.9±0.7 log_10_ and 1.7±1.0 log_10_ genome copies/10^6^ B cells during AIM and CONV, respectively.

### Deep sequencing reveals a high level of TCR diversity and epitope-specific features that drive selection of the TCR repertoire

To examine features that drive selection of YVL-and GLC-specific TCRs in AIM and CONV, deep sequencing of TCRα and β repertoires was conducted directly *ex vivo* on tetramer-sorted CD8 T cells at both time points (**Fig 1–2, S1–2, Table S2**). The characteristics of the TCR repertoires for each of 3 donors were elucidated by systematically analyzing preferential VA or VB usage hierarchy as presented in pie charts, CDR3 length analyses, VJ pairing by circos plots, and dominant CDR3 motif; the latter determines if there was enrichment of particular aa residues at specific sites important for ligand interaction. Enrichment for certain characteristics would suggest that these features are important for pMHC interaction and selection of the TCR repertoire^9,27,32–35^. YVL- and GLC-specific CD8 TCR repertoires in AIM demonstrated inter-individual differences, and were highly diverse; the mean (±SD) number of unique clonotypes were not significantly different in CONV (YVL: AIM TCRα 5548±2118, TCRβ 9038±3208; CONV TCRα 2981±940, TCRβ 3899±626; GLC: AIM TCRα 4202±1782, TCRβ 4540±1028; CONV TCRα 3982±848, TCRβ 4964±1139). However, there was a pronounced bias toward particular VA and VB gene family usage, specific to each epitope, that was maintained from AIM into CONV.

**Figure 1:**
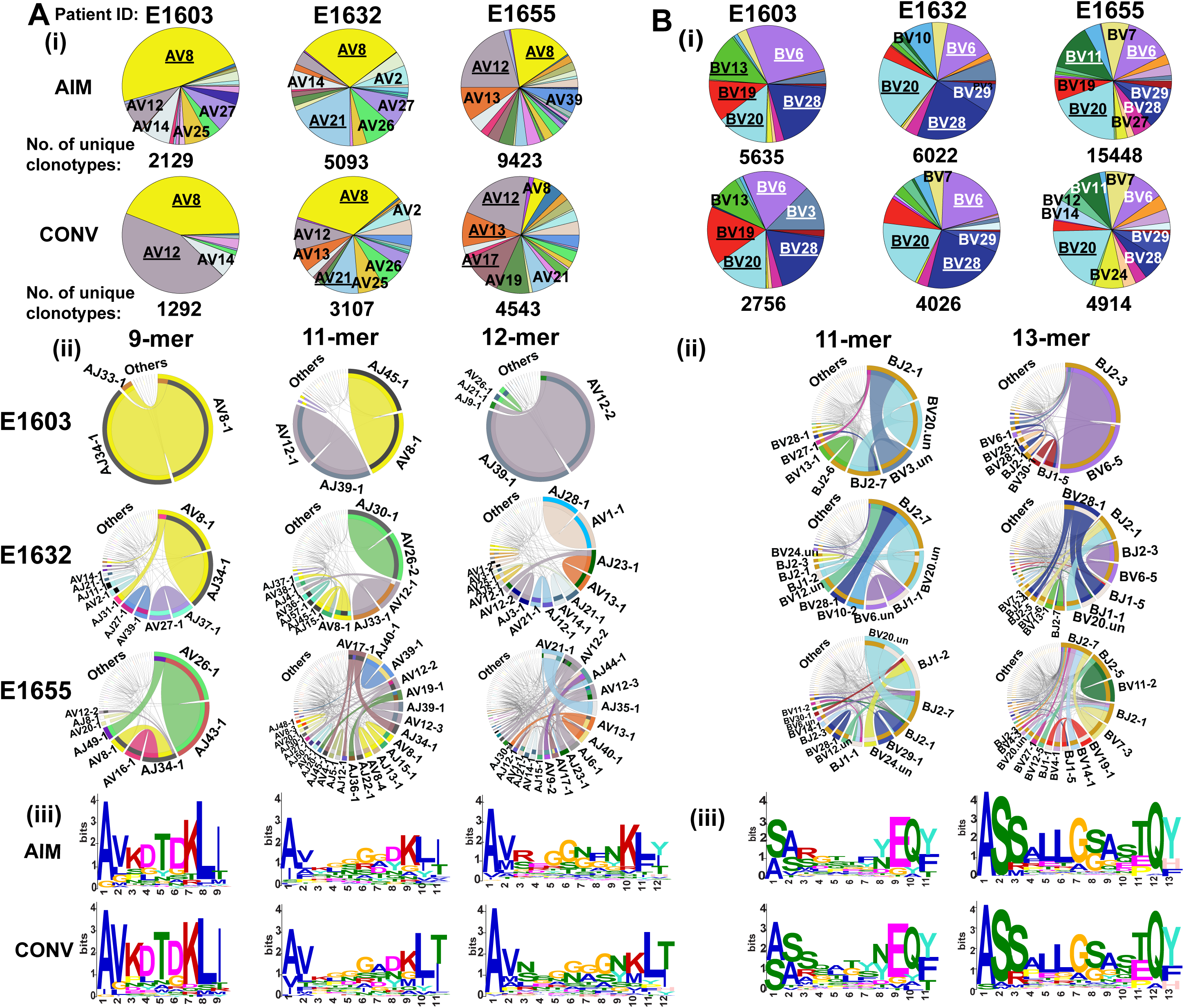
9-mer TCRα AV8.1-VKDTDK-AJ34 drives the selection of YVL-specific CD8 T-cells in AIM and CONV. HLA-A2/YVL-specific TCRAV (A) and TCRBV (B) repertoires were analyzed for 3 AIM donors (E1603, E1632, E1655) during the acute (within two weeks of onset of symptoms; primary response) and convalescent (6 months later; memory response) phase of EBV infection. Frequency of each TRAV (A) and TRBV (B) in total HLA-A2/YVL-specific TCR-repertoire is shown in pie charts (i). The pie plots are labeled with gene families having a frequency ≥10% (dominant, underlined) or between 5% and 10% (subdominant; not underlined). The total numbers of unique clonotypes in each donor is shown below the pie charts. There is consistent usage of AV8 and AV12 genes in all 3 donors in both the primary and memory phases. (ii) circos plots depicting V-J gene pairing and (iii) CDR3 motif analysis for the clonotypes with the two most dominant CDR3 lengths. Circos plots are only shown for the memory phase (AIM circos plots in Fig S1A). The frequencies of V-J combinations are displayed in circos plots, with frequency of each V or J cassette represented by its arc length and that of the V-J cassette combination by the width of the arc. “.un” denotes V families where the exact gene names were unknown.

**Figure 2:**
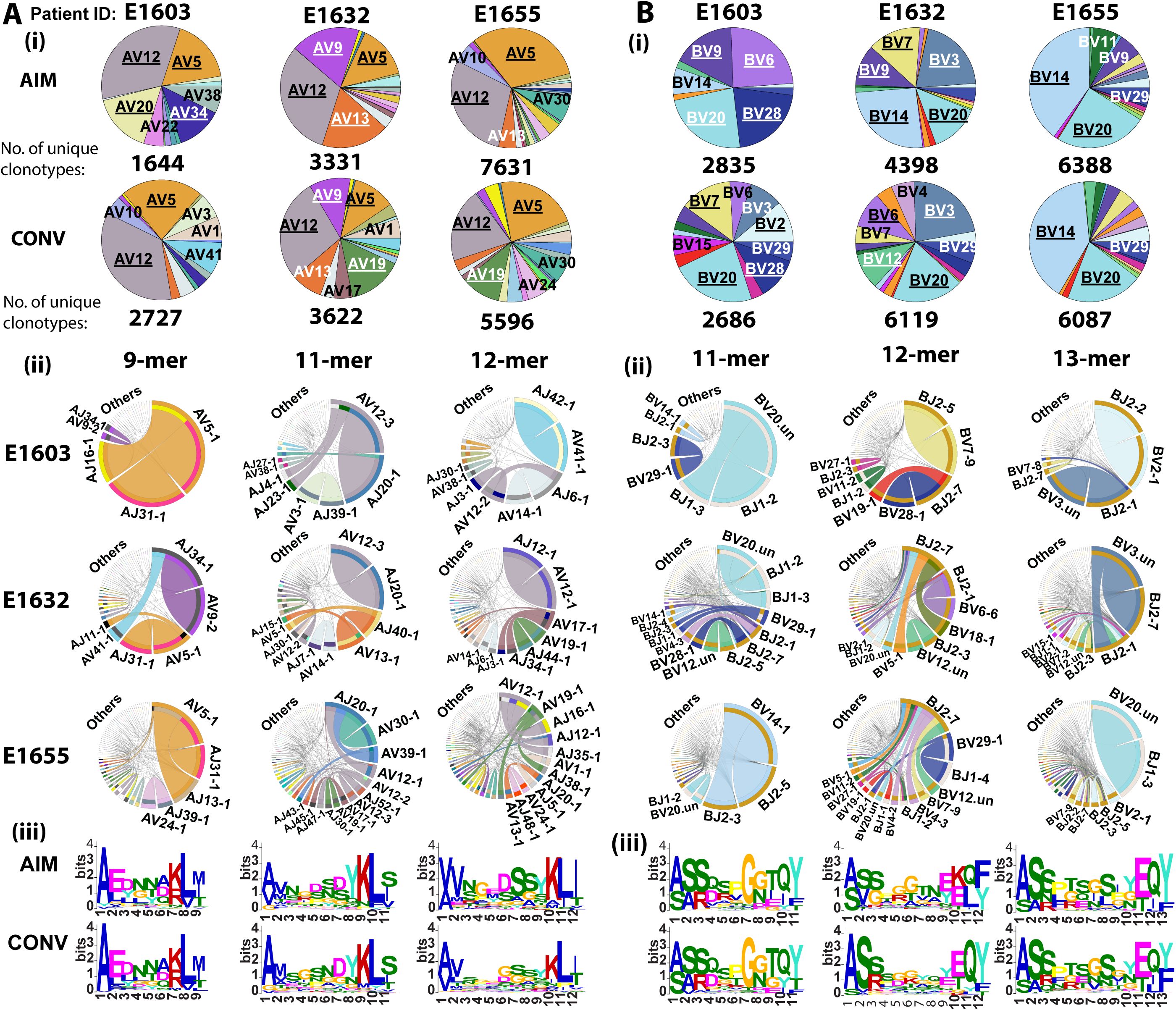
TCRα, AV5-EDNNA-AJ31, and TCRβ, BV14-SQSPGG-BJ2 and BV20-SARD-BJ1, clones are dominant selection factors for GLC-specific CD8 T-cells in AIM and CONV. HLA-A2/GLC-specific TCRAV (A) and TCRBV (B) repertoires were analyzed for 3 AIM donors (E1603, E1632, E1655) during the acute (within two weeks of onset of symptoms; primary response) and convalescent (6 months later; memory response) phase of EBV infection. Frequency of each TRAV (A) and TRBV (B) in total HLA-A2/GLC-specific TCR-repertoire is shown in pie charts (i). The pie plots are labeled with gene families having a frequency ≥10% (dominant, underlined) or between 5% and 10% (subdominant; not underlined). The total numbers of unique clonotypes in each donor is shown below the pie charts. There is consistent usage of AV5, AV12 and BV20 genes in all 3 donors in both the primary and memory phases. Otherwise, there is a high degree of variability in other AV and BV usage between donors and times. (ii) circos plots depicting V-J gene pairing and (iii) CDR3 motif analysis for the clonotypes with the two most dominant CDR3 lengths. Circos plots are only shown for the memory phase. (AIM circos plots in Fig S1B). “.un” denotes V families where the exact gene names were unknown.

### The 9-mer TCRα AV8.1-VKDTDK-AJ34 drives selection of YVL-specific CD8 T-cells

The YVL-specific TCR AV repertoire was focused on one dominant family, AV8, used by all donors in AIM and CONV (**Fig 1Ai, S1AAi**). Similar strong selection bias was not observed in YVL-specific BV usage; there was a great deal of inter-individual variation and preferential usage of multiple families, including BV6, BV20, BV28, BV29 (Fig 1Bi, S1ABi). However, in CONV some V gene families that dominated in AIM became extinct or subdominant, or new dominant genes emerged. For example, in CONV, E1655’s YVL TCR repertoire, AV8, BV6, and BV11 became subdominant while BV3 in E1603 and AV17 in E1655 became dominant (**Fig 1Ai, Bi**). In AIM, CDR3α length distribution varied from 9 to 14 aa; 9- and 11-mers represented 50% of all clonotypes (**Fig S1AAii**). Circos plot analysis of the 9-mer clonotypes showed that the dominant AV8.1 gene almost exclusively paired with AJ34 (**Fig S1AAiii**). CDR3α motif analysis revealed a pronounced motif, “VKDTDK”, in these shorter 9-mer clonotypes, representing 13.8%±5.6 of the total CD8 T-cell response during acute AIM **(Fig 1Aiii, S1AAiv, Table S3A)**; 87%±1.7 of the clonotypes using this motif were AV8.1 and 92%±1.7 were AJ34. Interestingly, this motif was present in multiple other AV and AJ pairs, including AV12, AV21 and AV3 among the most common. The fact that a dominant AV8.1 response obligately paired to AJ34 containing a highly conserved motif was observed in all donors from AIM through CONV, suggests that 9-mer AV8.1-VKDTDK-AJ34 expressing clones were highly selected by HLA-A2/YVL. The YVL CDR3β length varied, with 11- and 13-mer representing 60% of clonotypes. There was a preferential usage of BV20-BJ2.7 pairing within the 11-mer response **(Fig 1Bii, S1ABiii)** without a CDR3β motif **(Fig 1Biii, S1ABiv)**, highlighting a great degree of diversity in the aa composition. Within the 13-mer response **(Fig 1Biii,S1ABiv, Table S3B)**, the CDR3β motif, “LLGG”, was commonly used. This motif arose predominantly from BV28-BJ1 pairing (donor E1603, E1632) or BV6-BJ2 pairing (E1655). Clonotypes with this motif were only a minor part of overall responses in 2 donors (E1603, E1655), but composed 17.4% of the total YVL TCR BV repertoire in E1632. Altogether, these results suggest that the 9-mer AV8.1-VKDTDK-AJ34 expressing clones were highly preferentially selected by YVL ligand during AIM and CONV and that this AV chain could pair with multiple different BV chains.

### AV5-EDNNA-AJ31, BV14-SQSPGG-BJ2, and BV20-SARD-BJ1 are dominant selection factors for GLC-specific CD8 T-cells

The pattern of GLC-specific AV and BV usage had clear preference for particular gene families, consistent with prior reports^36,37^, that was maintained from AIM into CONV. We observed apparent preferential use of public AV5, 12, and BV20, 14, 9, 28, 29 families **(Fig 2Ai, Bi, S1BAi, Bi)**. There were differences in preferential usage of CDR3 lengths between the two epitopes. For instance, the AIM YVL-specific repertoire used more of the shorter 10-mer CDR3β than GLC in both AIM and CONV (**Fig 3Aii**). Like YVL, there were some individual changes in the transition into CONV with AV20 becoming extinct in E1603; AV19 emerging as dominant in E1632 (**Fig 2Ai**). In AIM, clonotypes with 9- and 11-mer CDR3α lengths represented 70% of the total response, while clonotypes with longer 11- and 13-mer CDR3β lengths represented 65% of total response (**Fig S1BAii, S1BBii)**. Circos plot analysis of the 9-mer CDR3α length clonotypes revealed a conserved and dominant AV5-AJ31 pairing in all 3 donors **(Fig S1BAiii)**. A prominent motif, “EDNNA”, was identified within 9-mer clonotypes, of which 85%±11 were associated with AV5-AJ31 (**Fig 2Aiii, S1BAiv, Table S3C**). This AV CDR3α motif was used by only 2.8%±1.7 of all clonotypes recognizing GLC in the 3 donors. The AV12 family dominated the 11-mer response, including AV12.1, AV12.2 and AV12.3 usage pairing with multiple different AJ genes depending on the donor. The 11-mer CDR3β BV14-BJ2 pairing exhibited a conserved, previously reported public motif, “SQSPGG”^38^, which represented 26% and 40% of the total GLC-specific response in donors E1632 and E1655 in AIM, respectively **(Fig S1BBii-iv, Table S3D)**. Within the CDR3β 13-mer responses, a conserved BV20-BJ1 pairing, including the previously reported public motif, “SARD”, was used by all 3 donors, and represented 11%±6 of the total GLC-specific response **(Fig 2Biii, S1BBii-iv, Table S3E)**. Within the 13-mer CDR3β response, there was also a consensus motif, “SPTSG” present in all 3 donors, which was used by multiple different BV families, which represented 20% and 2% of the total response in donors E1632 and E1655, respectively in AIM **(Fig 2Bii-iv, Table S3E)**. These data suggest that, in contrast to YVL whose TCR-repertoire selection was primarily driven by AV chain usage, the selection of GLC-specific TCR-repertoire in AIM was driven by a combination of both AV and BV chain usage.

**Figure 3:**
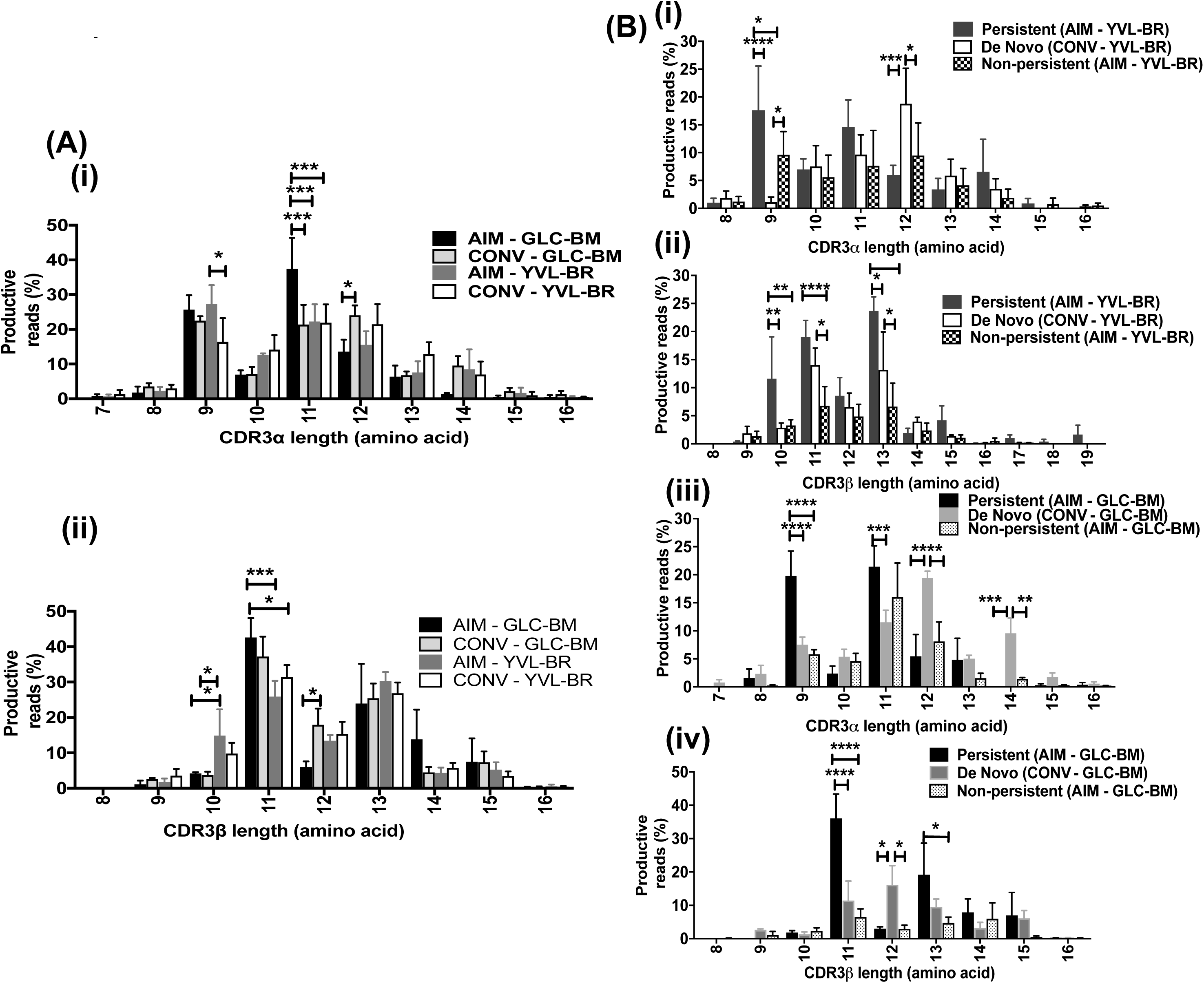
CDR3α and β length distribution of GLC- and YVL-specific CD8 T-cells in AIM and CONV (A) and in persistent, *de novo* and non-persistent clonotypes differ (B). (A) The mean CDR3 length distribution of the 3 EBV-infected patients’ TCR repertoire was analyzed by deep-sequencing of tetramer sorted cells during AIM and CONV. (B) The TCR repertoires were analyzed also after dividing each patients samples into 3 groups, those that persist from AIM into CONV and those which do not as well as *de novo* clonotypes arising in CONV. Data was analyzed by two-way ANOVA multivariant analysis, * p<0.05, ** p<0.01, *** p<0.001, **** p<0.0001. Error bars are SEM.

### CDR3 lengths are a major factor in the selection of the GLC- and YVL-specific TCR repertoires

The changing pattern of the epitope-specific TCR AV or BV family use from AIM to CONV for both epitope specific responses was also reflected by some changes in dominant CDR3 lengths (**Fig 3**). For example, the frequencies of the longer GLC-specific 12-mer CDR3 α and β clonotypes significantly increased from 13.6±6% and 6±2.8%, respectively, in AIM to 24±5% and 17.9±8%, respectively, in CONV; while use of the shorter 11-mer CDR3α decreased (**Fig 3Ai-ii)**. Within the YVL response usage of the shorter 9-mer CDR3α also decreased. Despite these changes, the factors that were identified in AIM to drive the selection of YVL- or GLC-specific CD8 T-cells were conserved in CONV, retaining the same dominant V gene families and CDR3 motifs indicating the strength of these TCR features in driving selection of the repertoire (**Fig 1–2, Table S3**).

To address whether clonotypes that persisted into memory show similar characteristics to those that dominate in acute infection, YVL and GLC TCRα/β repertoires were compared between AIM and CONV. Each unique TCRα or TCRβ clonotype (defined as a unique DNA rearrangement) elicited during AIM that was also detected during CONV was defined as a “persistent” clonotype. Clonotypes were regarded as “non-persistent” or “de novo” if they were present only during AIM or CONV, respectively. A high level of TCR diversity was maintained from AIM to CONV; however, the overlap between the number of unique clonotypes detected during AIM and CONV was small (**Fig 4Ai, Bi**). Only a small fraction, 6.6±2.2 - 9.1±4.2% of the TCRα/β YVL- and GLC-specific unique clonotypes, respectively, present in AIM were maintained at 8.7±4.9 - 18.5±5.6% during CONV. However, they comprised 57.5±26.2 - 75.5±12% of the total response when including their frequency (sequence reads) in AIM and 35.8±10.2 - 55.8±13.4% in CONV. While the clonotypic composition of GLC-and YVL-specific CD8 T-cells changed over the course of primary infection, dominant TCR clonotypes detected during AIM tended to persist and dominated in CONV. Altogether, these data indicate that persistent clonotypes made up only a small percentage of unique clonotypes but were highly expanded in AIM and CONV. Surprisingly, the vast majority (91%) of unique clonotypes completely disappeared following AIM and were replaced with *de novo* clonotypes in CONV.

**Figure 4:**
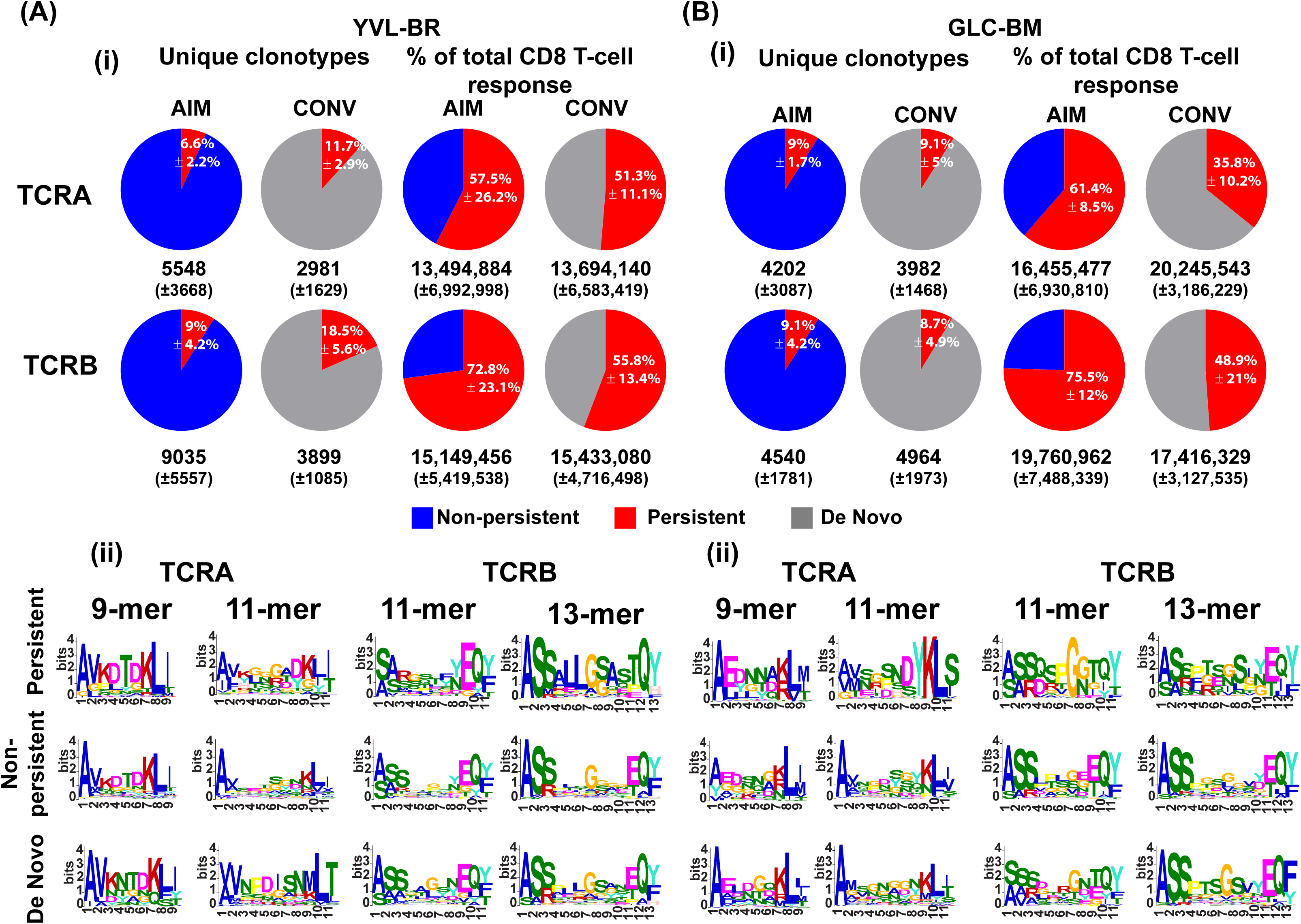
YVL- and GLC-specific structure drives selection of dominant persistent clonotypes while diversity is replaced but maintained. **(i)** Clonotypes that persist from the acute phase into memory represent only 6–18% of the unique clonotypes, but contribute to 35–75% of the total CD8 T-cell response. The highly diverse non-persistent clonotypes are replaced by new (*de novo*) highly diverse clonotypes, which were not present in the acute response. **(ii)** In both GLC and YVL responses, persistent clonotypes have characteristic CDR3 motifs that are distinct from non-persistent clonotypes, which tend to be very diverse. The *de novo* clonotypes appear to have new and unique CDR3 motifs. HLA-A2/YVL- (A) and GLC- (B) specific TCRAV and TCRBV repertoires were analyzed for 3 AIM donors (E1603, E1632, E1655) during the acute (within two weeks of onset of symptoms; primary response) and convalescent (6 months later; memory response) phase of EBV infection. The average frequency of unique clonotypes that persist into the memory phase (TRAV and TRBV) in total HLA-A2/YVL-specific (A) and GLC-specific (B) TCR-repertoire is shown (i). The average numbers (±SD) of unique clonotypes from the 3 donors are shown below the pie charts. Also shown in pie charts are the percentages of these unique clonotypes which persist into memory phase (TRAV and TRBV) and their percentage in the total CD8 T-cell response in total HLA-A2/YVL-specific (A) and GLC-specific (B) TCR-repertoire is shown (i). The average numbers (±SD) of sequence reads is shown below the pie charts.

The TCR repertoire of persistent and non-persistent clonotypes in AIM, and *de novo* clonotypes in CONV, were examined in order to identify selection factors that governed TCR persistence. Persistent YVL TCR clonotypes maintained the major selection factors that were identified in AIM (**Fig 4Aii, S2A-B)**. Although some features were maintained in all 3 TCR subsets, there were significant structural differences in these repertoires. Specifically, persistent clonotypes used significantly more of the shorter 9-mer CDR3α and more of 10-, 11-, and 12-mer CDR3β than the non-persistent. In contrast, the *de novo* clonotypes favored 12-mer CDR3α and 11-mer CDR3β length (**Fig 3Bi-ii**).

Non-persistent CDR3α clonotypes used AV8.1 but it was paired with many more AJ gene families **(Fig S2A**). Moreover, AV8.1-VKDTDK-AJ34 clonotypes, which were present in 42±20% or 19±11% of all persistent clonotypes during AIM or CONV, respectively, were present in the non-persistent response at a much lower mean frequency (6±1%; **Fig 4Aii, Table S4A-B)**. The clonal composition of the CDR3β non-persistent response varied greatly in BV usage between donors and lacked identifiable motifs, suggesting that for YVL clones expressing AV8.1-VKDTDK-AJ34 to persist, there are some preferential if not obvious TCRβ motifs.

For *de novo* clonotypes, new selection factors appeared that may relate to either a decrease in antigen expression or a change in antigen-expressing cells over the course of persistent infection. For instance, in the YVL 9-mer *de novo* clonotypes, the selection factor AV8.1-AJ34 was maintained in 2/3 donors and a new modified motif, VKNTDK was identified (**Fig 4Aii, S2Ai**). The *de novo* 11-mer CDR3α response had increased usage of AV12 in all 3 donors (**Fig S2Aii**). In *de novo* BV clonotypes, the pattern of BV-BJ usage changed compared to that observed in AIM. Similarly, *de novo* 13-mer CDR3β clonotypes were also totally different with usage of a new motif, SALLGX, in 2/3 donors **(Table S4C)**.

Within the GLC TCR repertoire, significant changes in TCR CDR3 length (**Fig 3Biii-iv**) were observed over time. The persistent clonotypes preferentially used 9- and 11-mer CDR3α while *de novo* preferred longer 12- and 14-mer lengths. The persistent clonotypes also preferentially used 11- and 13-mer CDR3β, while *de novo* preferred 12-mer lengths. The persistent GLC TCRα clonotypes maintained the major selection criteria that were identified in AIM with the 9-mer EDNNA motif, which strongly associated with AV5–1-AJ31, being present in a mean 5±3.7% or 10±8.6% of all persistent clonotypes during AIM or CONV, respectively, in all 3 donors (**Fig 4Bii, Table S4D)**. The fact that clonotypes using this motif were not present in non-persistent clonotypes suggests that this motif, and not just the gene family, may be important in determining persistence of GLC-specific clonotypes. The persistent GLC-repertoire also maintained the major selection criteria that were identified in AIM, with the 11-mer SARD motif that strongly associated with BV20.1-BJ1 being present in a mean 16±9.9% or 24±13.7% of all persistent clonotypes during AIM or CONV, respectively in all 3 donors. Two of the donors had the 11-mer SQSPGG motif in a mean 40±8% and 30±25% of all persistent clonotypes during AIM or CONV, respectively. Only the SARD motif clonotypes appeared in non-persistent BV clonotypes during AIM but at a lower mean frequency of 3±1% (**Table S4E)**. The *de novo* clonotype selection appeared to be driven by different factors than the persistent. Although there was much greater diversity and more variation between patients in *de novo* clonotypes (each donor is private) with recruitment of private AV families such as AV41 or AV24 in E1632 and E1655, there was still a preferential usage by 2/3 donors of AV5.1 (**Fig S3Ai)** and the appearance in 2/3 donors of a new 11-mer CDR3α motif “ELDGQ”, which associated with AV5.1-AJ16.1 (**Fig 4Bii, Table S4F)**. *De novo* clonotypes were also diverse and private using uncommon BV like BV7, BV3 but also using common BV families such as BV20 (**Fig S3B)** expressing the SARD motif in 5%±2.9 of *de novo* clonotypes(**Fig 4Bii)**.

In conclusion, the persistent clonotypes made up the vast majority of the AIM and CONV responses. For the most part, the non-persistent clonotypes did not have a motif despite the observation that some of them used a public AV or BV; this suggests that one of the strongest selection factors was the CDR3 motif. Additionally, the fact that persistent clonotypes retained features that were identified in AIM further supports their validity. Altogether, these results suggest that YVL- and GLC-specific structure drives selection of dominant persistent clonotypes while diversity is replaced but maintained.

### Single-cell paired TCRαβ sequencing confirms that selection of the YVL-specific repertoire is mainly driven by TCRα while GLC-specific repertoire selection is driven by unique combinations of TCRα/β

Single-cell TCR sequencing allows examination of TCRα/β pairing and gives even more accurate information on the structural constraints of pMHC/TCR interactions although it provides only limited information on TCR repertoire diversity. Circos plot analysis of AV-BV pairing of YVL-specific TCRs in AIM and CONV showed that the TCRα repertoire was more restricted while TCRβ was diverse. As suggested by the deep sequencing data, AV8.1 paired with a multitude of BV genes (most pronounced in E1651 and E1655) (**Fig 5A, C Table S5A)**. In GLC responses, there was less diversity and a conserved public AV5.1-BV20.1 pairing was present in all donors (**Fig 5B, D Table S5B)**.

**Figure 5:**
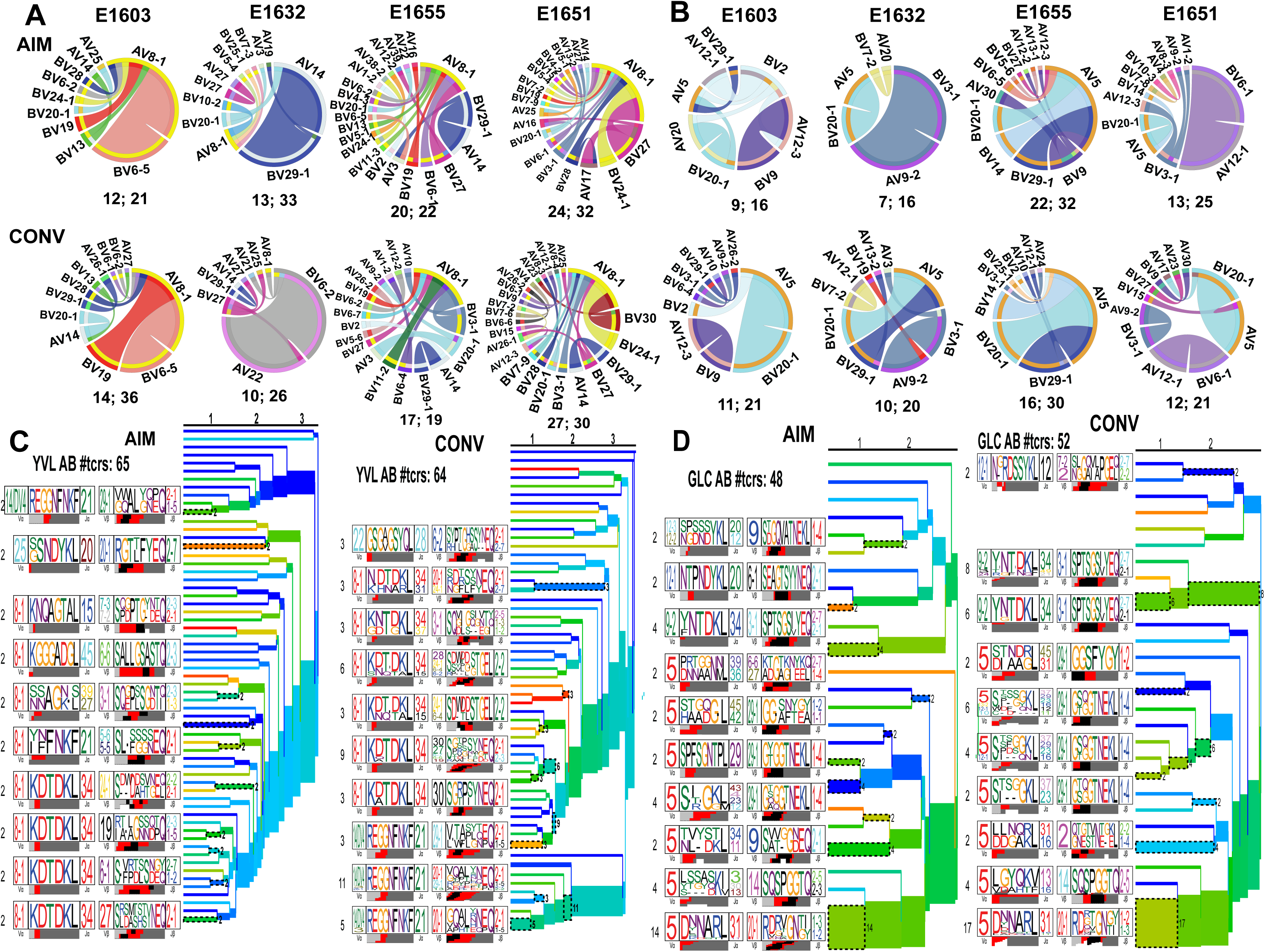
Patterns of AV-BV pairings by YVL (A) and GLC (B) specific CD8 T-cells as revealed by single-cell TCRαβ sequencing. The frequencies of AV-BV combinations in four AIM donors for YVL- (A) and GLC-specific (B) TCRαβ repertoires are displayed in circos plots, with frequency of each AV or BV cassette represented by its arc length and that of the AV-BV cassette combination by the width of the joining ribbon. The numbers of unique and productive paired TCRαβ clonotypes as well as the total numbers of sequences for each donor are shown below the pie charts (# of unique TCRαβ clonotypes; total # of sequences). Hierarchal clustering of TCRs highlights the structural features required for interaction with pMHC of paired TCRα/β: TCRαβ clustering along with corresponding TCR logos for (C) YVL-and (D) GLC-specific CD8 T-cell responses in AIM and CONV. Number on the branches and next to TCR logos depicts number of TCRs contributing to the cluster. Color of the branches indicates the TCR probability generation scores (based on Dash *et al.^9^*).

TCRα/β pairing was further characterized by examining the pattern of gene segment usage by ribbon plots, correlating gene usage within and across TCRα/β, quantifying preferential gene usage and CDR3 motif*^9^ (***Fig 6**). The results were combined from the 4 individuals by time point and epitope specificity. In AIM and CONV, the AV YVL-repertoire was focused on AV8.1, while the BV repertoire was highly diverse (**Fig 6A)**. The public and dominant AV8.1 used by YVL-specific TCRs preferentially combined with AJ34 and yet did not display any strong preference for any particular BV, and BJ (**Fig 6A)**. In contrast, both the TCRα/β GLC-repertoires were more focused and characterized by a very dominant gene combination involving AV5.1/AJ31/BV20.1/BJ1 (**Fig 6A)**. Jensen-Shannon divergence analysis used to quantify the total magnitude of gene preference confirmed that while GLC-specific TCRs showed a strong preferential usage of particular AV and BV, YVL-specific TCRs only had a strong preference for particular AV **(Fig 6B)**. Quantification of the degree of gene usage co-variation between pairs of segments using the adjusted mutual information score revealed that GLC-specific TCRs displayed a high stringency, every TCR gene combination except for two in AIM (AV-AJ and AV-BJ pairings) and one in CONV (AV-BJ) were important for selection into the repertoire **(Fig 6C)**. In contrast, no obvious gene associations emerged for YVL-specific TCR in AIM, whereas only the AV-AJ association emerged as important in CONV for selection into the TCR repertoire **(Fig 6C)**. CDR3α and β motif analysis at the single cell level demonstrated similar findings to those observed from deep-sequencing (**Fig 6D**). In summary, single-cell analysis complemented deep-sequencing analysis by providing direct evidence that the stringency for selection into either EBV-epitope specific TCR-repertoire differed greatly. Only very particular TCRAV (which could pair with multiple different TCRBV) was required to be selected into the YVL-specific TCR-repertoire, whereas very particular pairing of TCRAV and BV, with just the right J regions are required to be selected into GLC-specific TCR-repertoire.

**Figure 6.**
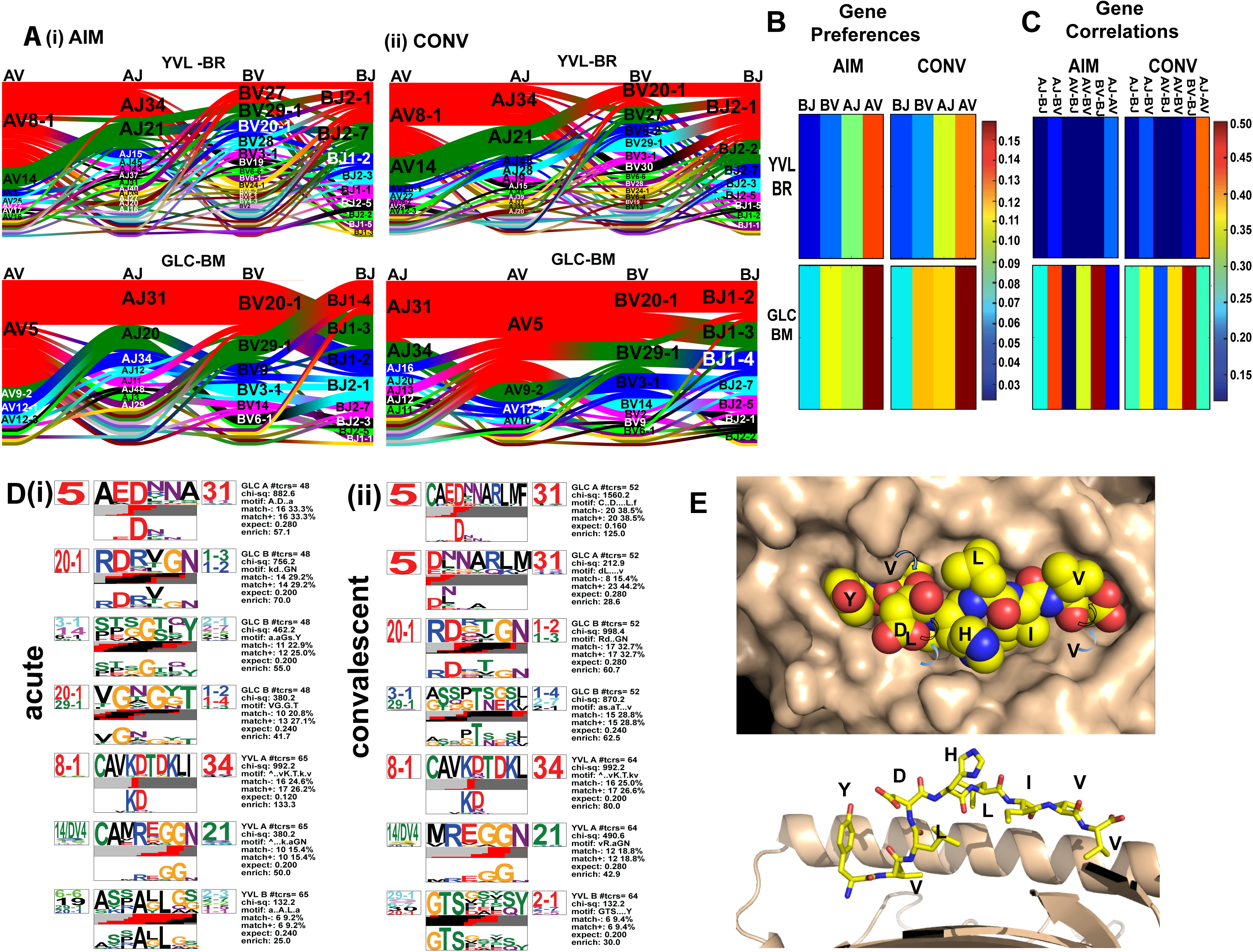
Single-cell paired TCRαβ sequencing provides further evidence that the selection of YVL-specific repertoire is mainly driven by TCRα while GLC-specific repertoire selection is driven by unique combinations of TCRα/β. (A) Gene segment usage and gene–gene pairing landscapes are illustrated using four vertical stacks (one for each V and J segment) connected by curved paths whose thickness is proportional to the number of TCR clones with the respective gene pairing (each panel is labeled with the four gene segments atop their respective color stacks and the epitope identifier in the top middle). Genes are colored by frequency within the repertoire which begins red (most frequent), green (second most frequent), blue, cyan, magenta, and black. (B), Jensen–Shannon divergence between the observed gene frequency distributions and background frequencies, normalized by the mean Shannon entropy of the two distributions (higher values reflect stronger gene preferences). (C), Adjusted mutual information of gene usage correlations between regions (higher values indicate more strongly covarying gene usage). (D) Single cell CDR3α and β motif analyses. Analyses in A-D based on Dash *et al*^9^. (E) Crystal structure of YVL pMHC*. Top*, top view (or TCR-view) of HLA-A2/YVL complex, with peptides atoms shown as spheres and HLA-A2 surface in tan. Peptide sequence indicated in single letter code. *Bottom*, cut-away side view of HLA-A2/YVL complex with peptide in stick model and HLA-A2 as a ribbon. The alpha-2 helix in front of the peptide was removed for clarity.

### Structural features support the role of Vα in selection of the YVL-specific TCR repertoire

We determined the crystal structure of the HLA-A2/YVL complex in order to identify specific structural features that might be associated with the dominant TRAV and TRAJ usage in the YVL-specific TCRs (**Fig 6E**). Structure determination was complicated initially by the relatively low resolution (**Table S6**) and high degree of translational pseudosymmetry (**Fig S4A-D**) that characterized the HLA-A2/YVL crystals (see Methods), but ultimately clear and continuous electron density extending the full length of the peptide allowed confident location of all peptide atoms (**Fig S4E-F**). In the HLA-A2 complex, the nonameric YVL peptide is bound in conventional orientation, with the amino terminus and side chains of valine at position 2 and valine at position 9 accommodated in pockets (A, B, and F, respectively) in the peptide binding site. Notably, the side chains of tyrosine at position 1, highly negative charged aspartic acid at position 4, and histidine at position 5 project away from the binding site, and compose a highly featured surface positioned for recognition by CDRα loops of a TCR bound in conventional orientation (**Fig 6E**). By contrast, the side chains of leucine at position 6, isoleucine at position 7, and valine at position 8, together with exposed main-chain atoms, present a rather non-descript generally hydrophobic surface positioned to interact with CDRβ loops. This structure would predict that positively charged Lys(K)/Arg(R) in the highly dominant YVL-specific TCRα motifs (**Fig 6D**) might be important for recognizing the asp(D) at position 5.

## Discussion

This is the first study to undertake a comprehensive longitudinal investigation of both the TCRα and β repertoires to two different epitope-specific CD8 T-cell responses in the context of a primary human EBV infection. Here, we show that each epitope drives the selection of highly diverse repertoires, that vary greatly between donors, but each epitope selects for distinct dominant clones with public features that persist into convalescence. These persistent clonotypes have distinct features specific to each antigen that appear to drive their selection; they account for only 9% of unique clonotypes, but predominate in acute infection and convalescence, accounting for 57%±4 of the total epitope-specific response. Surprisingly, the other 91% of highly diverse unique clonotypes disappear following AIM and are replaced in convalescence by equally diverse “*de-novo*” clonotypes (43% + 5% of the total response). The selection of unique public TCR repertoire features for each epitope in clones that dominate and persist suggest that these clones may be the best fit TCR to recognize each epitope. In contrast, the broad repertoire of unique clonotypes that are activated in AIM during high viral load and increased inflammation may not fit as well and perhaps do not receive a TCR signal that leads to survival into memory. Interestingly, 6 months after the initial infection, a completely new (*de novo)* and similarly diverse TCR repertoire has expanded. This may be due to continued antigenic exposure in persistent EBV infection. Earlier studies using similar techniques to study the influenza A (IAV) HLA-A*02:01-restricted IAV-M1_58–67_ and the cytomegalovirus (CMV)-pp65 epitope-specific memory responses showed a similar focused diversity structure in TCR repertoire, suggesting that this is a general principle of antigen-specific repertoire structure^27,28^. Altogether, these studies suggest that the pMHC structure drives selection of the particular public featured dominant clonotypes for each epitope. The broad fluctuating private repertoires show the resilience of memory repertoires and may lend plasticity to antigen recognition, perhaps assisting in early cross-reactive CD8 T-cell responses to heterologous new pathogens while at the same time potentially protecting against T-cell clonal loss and viral escape^39^.

These studies have also uncovered a role for TCRα-driven selection of the YVL-repertoire (Fig 6). To the best of our knowledge, this is the first study to describe a TCRα-driven selection of viral epitope-specific TCRs. AV8.1 was a public gene family, dominating the conserved 9-mer response, with an obligate pairing with AJ34, and a predominant CDR3 motif “VKDTDK”, representing 42% and 19% of the total persistent response in AIM and CONV, respectively. In contrast, the BV response was highly diverse without evidence of a strong selection factor, suggesting that AV8.1-VKDTDK-AJ34 could pair with multiple different BV and still successfully be selected by YVL/MHC as seen in the single cell paring data (**Table S5A)**. Unlike YVL, the selection of GLC-specific TCRs was driven by strong interactions with both TCRα/β, such as AV5.1-EDNNA-AJ31, BV14-SQSPGG-BJ2 and BV20.1-SARD-BJ1, including previously identified public features^36,37,40,41^. In a recent study comparing TCRα/β repertoires of various human and murine viral epitopes, none of the responses were primarily driven by interaction with TCRα alone; rather they were predominantly driven by strong interactions with TCRβ or a combination of TCRα/β^9^. Our YVL/MHC crystal structure data (Fig 6E) and structural modeling data^42^ shows a highly protuberant negatively charged aspartic acid (D) in position 4 of the peptide, in a location that TCRα would have to accommodate. The apparent preferential selection of AV8.1-VKDTDK-AJ34, which contains the positively charged K in position 2 would be consistent with this requirement. Future structural analysis would be important in confirming whether the TCRα contributes the majority of contacts with the pMHC, creating a large repertoire of different memory TCRβ that could potentially cross-react with other ligands such as IAV-M1, which predominantly interact with TCRBV^9,26,27^.

The deep sequencing results show a highly diverse TCR repertoire in each epitope-specific response with 1,292–15,448 and 1,644–7,631 unique clonotypes detected within the YVL and GLC-specific TCR-repertoires, respectively. Such diversity has been underappreciated for the GLC-specific TCR repertoire, with prior studies reporting an oligoclonal repertoire ^36,37,40^ of about 3–18 unique GLC-specific TCRβ clonotypes per individual. Despite this enormous diversity, there was considerable bias. Although the TCR repertoire was individualized (*i.e*., each donor studied had a unique TCR-repertoire), there was prevalent and public usage of particular gene families such as AV8 within the YVL-specific responses and AV5, AV12 and BV14, BV20 within the GLC-specific populations.

Close examination of our data reveals several insights into the mechanisms of TCR selection and persistence. First, prior studies have revealed that selective use of particular gene families can be explained in part by the fact that the specificity of TCR for a pMHC is determined by contacts made between the germline-encoded regions within a V segment and the MHC^43,44.^ In fact, the CDR3 of the YVL AV8.1-VKDTDK-AJ34 dominant clonotype is derived from germline for VA and JA except for insertion of one aa, K, in the second position, which as described above, is likely required to accommodate the negatively charged aspartic acid in position 4 of the peptide. Second, it has been suggested that public TCRs represent clonotypes present at high frequency in the naïve precursor pool as a result of bias in the recombination machinery^45^ or convergent recombination of key contact sites^41,44,46,47.^ Using the convergence theory and counting the total number of DNA arrangements in our data that have resulted in a given amino acid sequence (for TCRα and TCRβ separately) for persistent and non-persistent clonotypes (pooling responses to both epitopes), the median (mean) number of DNA sequences coding for CDR3 of one TCRα in persistent clonotypes was 5 (36.6) vs 1 (4.2) in non-persistent (p<2.2E-16 Wilcoxon test) and for one TCRβ in persistent clonotypes was 5 (33.3) vs 1 (5.2) in non-persistent (p<2.2E-16 Wilcoxon test). We obtained similar results with this type of analysis showing that public vs private clonotypes used more convergent amino acids in the CDR3α/β for both epitopes. Thus, persistent clonotypes and public clonotypes have TCRs with amino acid sequences coded for by a much larger number of DNA sequences consistent with the convergent theory of common or public TCR selection. Finally, we have previously reported that TCR immunodominance patterns also seem to scale with number of specific interactions required between pMHC and TCR^27^. It would seem that TCRs that find simpler solutions to being generated and to recognizing antigen involving fewer specific amino acids are easier to evolve and come to dominate the memory pool.

Despite the loss of the vast majority of the initial pool of clones deployed during acute infection, clonotypic diversity was unaffected and remained high in memory as a result of the recruitment of a diverse pool of new clonotypes. In a murine model, adoptive transfer of epitope-specific CD8 T-cells of known BV family from a single virus-infected mouse to a naive mouse, followed by viral challenge, resulted in altered hierarchy of the clonotypes and the recruitment of new clonotypes, thus maintaining diversity ^48^. A highly diverse repertoire should allow resilience against loss of individual clonotypes with aging^49^ and against skewing of the response after infection with a cross-reactive pathogen^50,51^. The large number of clonotypes contributes to the overall memory T-cell pool, enhancing the opportunity for protective heterologous immunity now recognized to be an important aspect of immune maturation^52^. A large pool of TCR clonotypes could provide increased resistance to viral escape mutants common in persistent virus infections^39^. Different TCRs may also activate antigen-specific cell functions differently, leading to a more functionally heterogeneous pool of memory cells^53^.

In summary, our data revealed that apparent molecular constraints were associated with TCR selection and persistence in the context of controlled viral replication following primary EBV infection. They also show that TCRAV can play an equally important role to TCRBV in TCR selection against important immunodominant responses; thus, to understand the rules of TCR selection, both repertoires need to be studied. Further studies could elucidate which of the features of the epitope-specific CD8 TCR are associated with an effective response and control of EBV replication or disease.

## Supplemental Figure Legends

**Supplemental Figure S1: Dominant selection factors of YVL- and GLC-specific TCRαβ repertoires during AIM as assessed by deep sequencing**. The TCR repertoire was deconstructed by analyzing V family usage in pie chart format, CDR3 length analyses, VJ pairing by using circos plot analyses, and CDR3 aa motif analyses using Multiple MEME framework^54^. **(S1A) 9-mer TCR**α **AV8.1-VKDTDK-AJ34 drives the selection of YVL-specific CD8 T-cells in AIM**. HLA-A2/YVL-specific TCRAV (A) and TCRBV (B) repertoires were analyzed for 3 AIM donors (E1603, E1632, E1655) during the acute (within two weeks of onset of symptoms; primary response) phase of EBV infection. Frequency of each TRAV (A) and TRBV (B) in total HLA-A2/YVL-specific TCR-repertoire is shown in pie charts (i). The pie plots are labeled with gene families having a frequency ≥10% (dominant, underlined) or between 5% and 10% (subdominant; not underlined). The total numbers of unique clonotypes in each donor is shown below the pie charts. (ii) CDR3 length distribution along with (iii) circos plots depicting V-J gene pairing and (iv) CDR3 motif analysis for the clonotypes with the two most dominant CDR3 lengths. Genes are colored by V gene family with a fixed color sequence used throughout the manuscript. **(S1B) TCR**α**, AV5-EDNNA-AJ31, and TCR**β**, BV14-SQSPGG-BJ2 and BV20-SARD-BJ1, clones are dominant selection factors for GLC-specific CD8 T-cells during AIM**. HLA-A2/GLC-specific TCRAV (A) and TCRBV (B) repertoires were analyzed for 3 AIM donors (E1603, E1632, E1655) during the acute (within two weeks of onset of symptoms; primary response) phase of EBV infection. Frequency of each TRAV (A) and TRBV (B) in total HLA-A2/GLC-specific TCR-repertoire is shown in pie charts (i). The pie plots are labeled with gene families having a frequency ≥10% (dominant, underlined) or between 5% and 10% (subdominant; not underlined). The total numbers of unique clonotypes in each donor is shown below the pie charts. There is consistent usage of AV5 and AV12 genes in all 3 donors. There is consistent usage of BV20 in all 3 donors. Otherwise there is a high degree of variability in other AV and BV usage between donors. (ii) CDR3 length distribution along with (iii) circos plots depicting V-J gene pairing and (iv) motif analysis for the clonotypes with the two most dominant CDR3 lengths. The frequencies of V-J combinations are displayed in circos plots, with frequency of each V or J cassette represented by its arc length and that of the V-J cassette combination by the width of the arc. “.un” denotes V families where the exact gene names were unknown.

**Supplemental Figure S2: Patterns of V-J usage for persistent, non-persistent and *de novo* clonotypes (9- and 11-mer CDR3α (A) and 11- and 13-mer CDR3β (B)) of YVL-specific CD8 T-cell responses as obtained by deep-sequencing**. The frequencies of V-J combinations in three AIM donors for YVL-specific TCRαβ repertoires are displayed in circos plots, with frequency of each V or J cassette represented by its arc length and that of the V-J cassette combination by the width of the arc. “.un” denotes V families where the exact gene names were unknown.

## Materials and Methods

### Study population

Four individuals (E1603, E1632, E1655 and E1651) presenting with symptoms consistent with acute infectious mononucleosis (AIM) and laboratory studies consistent with primary infection (positive serum heterophile antibody and the detection of EBV viral capsid antigen (VCA)-specific IgM) were studied as described^26^. Blood samples were collected in heparinized tubes at clinical presentation with AIM symptoms (acute phase) and six months later (memory phase). PBMC were extracted by Ficoll-Paque density gradient media. The Institutional Review Board of the University of Massachusetts Medical School approved these studies, and all participants provided written informed consent.

### Flow cytometry and isolation of GLC- and YVL-specific CD8 T-cells

The percentages of peripheral blood antigen-specific CD8 T-cells were measured using flow cytometry analysis. Antibodies included: anti-CD3-FITC, anti-CD4-AF700 and anti-CD8-BV786, 7AAD and PE-conjugated HLA-A*02:01-peptide tetramers (BRLF-1: **<u>YVL</u>**DHLIVV; BMLF-1: **<u>GLC</u>**TLVAML). Tetramers were made and underwent quality assurance, as previously described^55^.

Total CD8 T-cells were enriched from PBMC by positive selection using MACS technology (Miltenyi Biotec, Auburn, CA) according to the manufacturer’s protocol. The cells were then stained with anti-CD3, anti-CD4, anti-CD8, 7AAD, and GLC- or YVL-tetramers. Single-cell (into 384-well PCR plates) or bulk sorting (FACSAria III, BD) of live CD3+, CD8+, and GLC- or YVL-tetramer+ cells were sorted by flow cytometry for subsequent TCR analysis.

### Analysis of TCRαβ CDR3 regions using deep sequencing

Deep sequencing was performed for all donors except E1651 due to a lack of sample availability. Sequences of CDR3 regions were identified according to the definition founded by the International ImMunoGeneTics collaboration. Deep sequencing data of TCRα and β repertoires were analyzed using ImmunoSEQ Analyzer versions: 2.0 and 3.0, which were provided by Adaptive Biotechnologies. Only productively (without stop codon) rearranged TCRα and TCRβ sequences were used for repertoire analyses, including sequence aa composition and gene-frequency analyses. The frequencies of *AV-AJ* and *BV-BJ* gene combinations were analyzed with subprograms of the ImmunoSEQ Analyzer software and further processed by Microsoft Excel.

#### Circos plots and motif analysis

The V and J gene segment combinations were illustrated as circos plots^56^ across different CDR3 aa sequence lengths. Motif analysis was performed using the Multiple EM for Motif Elicitation (MEME) framework^54^. Consensus motifs were acquired across different CDR3 lengths and statistics on those motifs were computed with an in-house program called motifSearch and available at http://github.com/thecodingdoc/motifSearch.

### Single-cell paired TCRαβ analysis of EBV-specific CD8 T-cells

To examine TCRα and TCRβ pairing relationships, we conducted an *ex vivo* single-cell analysis of the paired TCRαβ repertoire of YVL- and GLC-specific CD8 T-cells from PBMCs of the 3 donors we have discussed thus far and one additional donor (E1651) in AIM and CONV. Following single-cell sorting, amplification of paired CDR3α and CDR3β regions was performed as previously described^57^ by multiplex-nested reverse transcriptase PCR, followed by Sanger sequencing of TCRα and TCRβ products. Sequences (see Table S5) were analyzed according to the IMGT/V-QUEST web-based tool^58^.

#### Ribbon plots, gene correlations and gene preferences

An analytical tool developed by Dash *et al.^9^* was used to characterize patterns of gene segment usage by ribbon plots, correlate gene usage within a chain (for example, AV-AJ, BV-BJ) and across chains (for example, AV-BV, AV-BJ), and to quantify gene preference usage (the quantification was done by comparing the gene frequencies in our epitope-specific repertoires to those seen in a background set of publicly available non-epitope-selected repertoire using the Shannon diversity index). The analysis was done by combining productive single-cell TCRαβ sequences from the 4 donors at each time point and for each epitope. A total of 65 and 64 (YVL; AIM and CONV) and 48–52 (GLC; AIM and CONV) productive paired TCRαβ sequences were generated.

### EBV DNA quantitation in B cells

B cells were purified from whole blood using the RosetteSep human B-cell enrichment cocktail according to the manufacturer’s recommendations (StemCell Technologies, Vancouver BC, Canada). Cellular DNA was extracted using QIAGEN DNeasy Blood & Tissue Kit (Valencia, CA). Each DNA sample was diluted to 5ng/ul and the Roche LightCycler EBV Quantitation Kit (Roche Diagnostics, Indianapolis, IN) was used to quantify EBV DNA copy number in the samples as recommended by the manufacturer. Reactions were run in duplicate. B cell counts in each sample were determined using a previously described PCR assay to quantify the copy number of the gene encoding CCR5 (two copies per diploid cell)^59^. Samples were normalized to B cell counts and EBV DNA copy number was calculated as DNA copy per 10^6^ B cells.

### Soluble HLA-A2/YVL protein production and crystallization

Soluble HLA-A2/YVL complexes were prepared by folding urea-solubilized bacterially-expressed inclusion bodies of HLA-A2 heavy chain and human L2-microglobin in the presence of 5mg/L synthetic YVL peptide essentially as described^60^, followed by concentration and buffer exchange into 10mM Tris-Cl (pH 8.0) using a tangential flow concentrator. Folded HLA-A2/peptide complexes were isolated from the buffer-exchanged folding mixture by a series of chromatography steps consisting of Hitrap Q and Mono Q ion exchange and S-200 gel-filtration columns (GE healthcare). Crystals were grown from purified HLA-A2/YVL by sitting drop vapor diffusion using 10.5% (w/v) PEG 4000, 35 mM Tris base/ HCl (pH 8.5), 70 mM Li_2_SO_4_. Crystals were briefly soaked in 1:1 mixture of saturated sucrose and reservoir buffer for cryoprotection and flash-frozen in liquid nitrogen and sent to LRL-CAT beamline at the Advanced Photon Source (Argonne, IL USA).

### HLA-A2/YVL Structure determination and refinement

Diffraction data extending to ~ 3.2Å collected from a single crystal were integrated and indexed using Mosflm^61^. Initially data were indexed in a C2 unit cell (189.2 × 49.7 × 291.6 Å, β=94.5°), with molecular replacement using Phaser^62^ identifying four copies per asymmetric unit of a HLA-A2 model^27^ with TFZ=22. However, refinement of this model stalled at Rfree = 0.42. Re-examination of the diffraction pattern identified a lattice of weak spots spaced between the stronger spots originally indexed (Supplemental Fig S4A), and the data were reindexed in a P2_1_ unit cell (189.9 × 100.2 × 292.4 Å, β=94.4°). The newly identified spots comprise the k=2n+1 and h+(k/2)=2n+1 sets (Supplemental Fig S4). Additional molecular replacement, symmetry considerations, and examination of composite-omit maps calculated using CCP4i^63^ identified 20 copies of HLA-A2 per asymmetric unit (Supplemental Fig S4B). The molecules are arranged in two layers viewed looking into the ac plane (Supplemental Fig S4C); slight differences can be observed between these layers, and between similarly oriented molecules within the same layer (Supplemental Fig S4D). These differences, along with four molecules (A,J,K,T) not identified in the C2 cell, explain the lower symmetry and strong translational pseudosymmetry that resulted in weak intensities for the k=2n+1 and h+(k/2)=2n+1 spots. After identification of the correct crystallographic and non-crystallographic symmetries, clear electron density covering all peptide atoms was observed in 20-fold averaged composite omit maps (Supplemental Fig S4E), and a model for the YVL peptide was built using Coot^64^. Refinement using Phenix^65^ proceeded smoothly despite the relatively low resolution when dihedral restraints to a higher-resolution reference model were provided. Models for the YVL peptide, which was not included in the reference model restraints, did not vary significantly between the non-crystallographically related copies (Supplemental Fig S4F). A paired refinement test^66^ confirms 3.3 Å as a suitable resolution cutoff. Final refinement statistics, shown in Supplemental Table S6, are within the range of other structures determined at this resolution in the Protein Data Bank. PyMOL (The PyMOL Molecular Graphics System, Version 2.0 Schrodinger, LLC) was used for graphical representation of molecular structures.

#### Statistics

GraphPad Prism version 7.0 for Mac OSX (GraphPad Software, La Jolla, CA) was used for all statistical analyses.

## Data Availability

Raw TCR deep sequencing data can be accessed at:

https://urldefense.proofpoint.com/v2/url?u=http-3A__clients.adaptivebiotech.com&d=DwQGaQ&c=WJBj9sUF1mbpVIAf3biu3CPHX4MeRjY_w4DerPlOmhQ&r=p6IL5ohbVyB2IGgNCmdbh-A5IMFqxKtq0WBpidjH1QE&m=hueuAoY7ZXzP9YMFmhGPKpu9iLorr5nv05XTqQklDuI&s=AdlhcrGwYqZ-QWYlQON5AJFRO88HSQe1qPUMaWRkQik&e=

The login information is as follows: Username

Password: gil-2018-review

## Acknowledgements

We are grateful to the study subjects for their participation. We thank George Corey and Jessica Conrad for obtaining clinical samples; Linda Lambrecht, Robin Brody, Anita Gautam, and Jennifer Henderson for expert technical assistance; Margaret McManus for data management and critical review of the manuscript; and Dr. Raymond Welsh for critical review of the manuscript. This research used resources of the Advanced Photon Source, a U.S. Department of Energy (DOE) Office of Science User Facility operated for the DOE Office of Science by Argonne National Laboratory under Contract No. DE-AC02–06CH11357. Use of the Lilly Research Laboratories Collaborative Access Team (LRL-CAT) beamline at Sector 31 of the Advanced Photon Source was provided by Eli Lilly Company, which operates the facility.

## Author Contributions

K.L. and L.K.S. obtained samples and conceived the study. L.K. and A.G. contributed to study design, and were primarily responsible for cell sorting and TCR sequencing. All authors contributed to data analyses. D.G. and R.C. performed all computational analyses. I.S. and L.S. performed and analyzed the data from the crystallographic studies. K.L., L.K.S., L.K., A.G., D.G., and L.J.S. assumed primary responsibility for writing the manuscript. All authors reviewed, provided substantive input, and approved of the final manuscript.

## References

1. Luzuriaga, K. & Sullivan, J.L. Infectious mononucleosis. N Engl J Med 362, 1993–2000 (2010).

2. Pender, M.P. & Burrows, S.R. Epstein-Barr virus and multiple sclerosis: potential opportunities for immunotherapy. Clin Transl Immunology 3, e27 (2014).

3. Crawford, D.H. Biology and disease associations of Epstein-Barr virus. Philos Trans R Soc Lond B Biol Sci 356, 461–473 (2001).

4. Loren, A.W., Porter, D.L., Stadtmauer, E.A. & Tsai, D.E. Post-transplant lymphoproliferative disorder: a review. Bone Marrow Transplant 31, 145–155 (2003).

5. Bollard, C.M., et al. Sustained complete responses in patients with lymphoma receiving autologous cytotoxic T lymphocytes targeting Epstein-Barr virus latent membrane proteins. J Clin Oncol 32, 798–808 (2014).

6. Bollard, C.M., Rooney, C.M. & Heslop, H.E. T-cell therapy in the treatment of post-transplant lymphoproliferative disease. Nat Rev Clin Oncol 9, 510–519 (2012).

7. Keymeulen, B., et al. Insulin needs after CD3-antibody therapy in new-onset type 1 diabetes. N Engl J Med 352, 2598–2608 (2005).

8. Pender, M.P., Csurhes, P.A., Burrows, J.M. & Burrows, S.R. Defective T-cell control of Epstein-Barr virus infection in multiple sclerosis. Clin Transl Immunology 6, e126 (2017).

9. Dash, P., et al. Quantifiable predictive features define epitope-specific T cell receptor repertoires. Nature 547, 89–93 (2017).

10. Glanville, J., et al. Identifying specificity groups in the T cell receptor repertoire. Nature 547, 94–98 (2017).

11. DeWitt, W.S., et al. Human T cell receptor occurrence patterns encode immune history, genetic background, and receptor specificity. Elife 7(2018).

12. Emerson, R.O., et al. Immunosequencing identifies signatures of cytomegalovirus exposure history and HLA-mediated effects on the T cell repertoire. Nat Genet 49, 659–665 (2017).

13. Attaf, M., Huseby, E. & Sewell, A.K. alphabeta T cell receptors as predictors of health and disease. Cell Mol Immunol 12, 391–399 (2015).

14. Attaf, M. & Sewell, A.K. Disease etiology and diagnosis by TCR repertoire analysis goes viral. Eur J Immunol 46, 2516–2519 (2016).

15. Cohen, J.I., Mocarski, E.S., Raab-Traub, N., Corey, L. & Nabel, G.J. The need and challenges for development of an Epstein-Barr virus vaccine. Vaccine 31 Suppl 2, B194–196 (2013).

16. Yanagi, Y., et al. A human T cell-specific cDNA clone encodes a protein having extensive homology to immunoglobulin chains. Nature 308, 145–149 (1984).

17. Zinkernagel, R.M. & Doherty, P.C. Restriction of in vitro T cell-mediated cytotoxicity in lymphocytic choriomeningitis within a syngeneic or semiallogeneic system. Nature 248, 701–702 (1974).

18. Hedrick, S.M., Cohen, D.I., Nielsen, E.A. & Davis, M.M. Isolation of cDNA clones encoding T cell-specific membrane-associated proteins. Nature 308, 149–153 (1984).

19. La Gruta, N.L., Gras, S., Daley, S.R., Thomas, P.G. & Rossjohn, J. Understanding the drivers of MHC restriction of T cell receptors. Nat Rev Immunol 18, 467–478 (2018).

20. Siu, G., et al. The human T cell antigen receptor is encoded by variable, diversity, and joining gene segments that rearrange to generate a complete V gene. Cell 37, 393–401 (1984).

21. Miles, J.J., Douek, D.C. & Price, D.A. Bias in the alphabeta T-cell repertoire: implications for disease pathogenesis and vaccination. Immunol Cell Biol 89, 375–387 (2011).

22. Nikolich-Zugich, J., Slifka, M.K. & Messaoudi, I. The many important facets of T-cell repertoire diversity. Nat Rev Immunol 4, 123–132 (2004).

23. Furman, D., et al. Cytomegalovirus infection enhances the immune response to influenza. Science translational medicine 7, 281ra243 (2015).

24. Kloverpris, H.N., et al. CD8+ TCR Bias and Immunodominance in HIV-1 Infection. Journal of immunology (Baltimore, Md. : 1950) 194, 5329–5345 (2015).

25. Watkin, L., et al. Potential of unique influenza A crossreactive memory CD8 TCR repertoire to protect against Epstein Barr virus (EBV) seroconversion. J Allergy Clin Immun **in press**(2017).

26. Aslan, N., et al. Severity of Acute Infectious Mononucleosis Correlates with Cross-Reactive Influenza CD8 T-Cell Receptor Repertoires. MBio 8(2017).

27. Song, I., et al. Broad TCR repertoire and diverse structural solutions for recognition of an immunodominant CD8+ T cell epitope. Nat Struct Mol Biol 24, 395–406 (2017).

28. Chen, G., et al. Sequence and Structural Analyses Reveal Distinct and Highly Diverse Human CD8+ TCR Repertoires to Immunodominant Viral Antigens. Cell Rep 19, 569–583 (2017).

29. Abdel-Hakeem, M.S., Boisvert, M., Bruneau, J., Soudeyns, H. & Shoukry, N.H. Selective expansion of high functional avidity memory CD8 T cell clonotypes during hepatitis C virus reinfection and clearance. PLoS Pathog 13, e1006191 (2017).

30. Klarenbeek, P.L., et al. Deep sequencing of antiviral T-cell responses to HCMV and EBV in humans reveals a stable repertoire that is maintained for many years. PLoS Pathog 8, e1002889 (2012).

31. Sant, S., et al. Single-Cell Approach to Influenza-Specific CD8(+) T Cell Receptor Repertoires Across Different Age Groups, Tissues, and Following Influenza Virus Infection. Front Immunol 9, 1453 (2018).

32. Yang, X., et al. Structural Basis for Clonal Diversity of the Public T Cell Response to a Dominant Human Cytomegalovirus Epitope. J Biol Chem 290, 29106–29119 (2015).

33. Pymm, P., et al. MHC-I peptides get out of the groove and enable a novel mechanism of HIV-1 escape. Nat Struct Mol Biol 24, 387–394 (2017).

34. Yang, X., Chen, G., Weng, N.P. & Mariuzza, R.A. Structural basis for clonal diversity of the human T-cell response to a dominant influenza virus epitope. J Biol Chem 292, 18618–18627 (2017).

35. Singh, N.K., et al. Emerging Concepts in TCR Specificity: Rationalizing and (Maybe) Predicting Outcomes. J Immunol 199, 2203–2213 (2017).

36. Annels, N.E., Callan, M.F., Tan, L. & Rickinson, A.B. Changing patterns of dominant TCR usage with maturation of an EBV-specific cytotoxic T cell response. J Immunol 165, 4831–4841 (2000).

37. Callan, M.F., et al. CD8(+) T-cell selection, function, and death in the primary immune response in vivo. J Clin Invest 106, 1251–1261 (2000).

38. Nguyen, T.H., et al. Maintenance of the EBV-specific CD8+ TCRalphabeta repertoire in immunosuppressed lung transplant recipients. Immunol Cell Biol 95, 77–86 (2017).

39. Wolfl, M., et al. Hepatitis C virus immune escape via exploitation of a hole in the T cell repertoire. J Immunol 181, 6435–6446 (2008).

40. Levitsky, V., de Campos-Lima, P.O., Frisan, T. & Masucci, M.G. The clonal composition of a peptide-specific oligoclonal CTL repertoire selected in response to persistent EBV infection is stable over time. J Immunol 161, 594–601 (1998).

41. Venturi, V., et al. TCR beta-chain sharing in human CD8+ T cell responses to cytomegalovirus and EBV. J Immunol 181, 7853–7862 (2008).

42. Antunes, D.A., et al. Interpreting T-Cell Cross-reactivity through Structure: Implications for TCR-Based Cancer Immunotherapy. Front Immunol 8, 1210 (2017).

43. Davis, M.M., McHeyzer-Williams, M. & Chien, Y.H. T-cell receptor V-region usage and antigen specificity. The cytochrome c model system. Ann N Y Acad Sci 756, 1–11 (1995).

44. Miles, J.J., et al. Genetic and structural basis for selection of a ubiquitous T cell receptor deployed in Epstein-Barr virus infection. PLoS Pathog 6, e1001198 (2010).

45. Yassai, M., et al. Naive T cell repertoire skewing in HLA-A2 individuals by a specialized rearrangement mechanism results in public memory clonotypes. J Immunol 186, 2970–2977 (2011).

46. Venturi, V., et al. Sharing of T cell receptors in antigen-specific responses is driven by convergent recombination. Proc Natl Acad Sci U S A 103, 18691–18696 (2006).

47. Venturi, V., Price, D.A., Douek, D.C. & Davenport, M.P. The molecular basis for public T-cell responses? Nat Rev Immunol 8, 231–238 (2008).

48. Cukalac, T., et al. Reproducible selection of high avidity CD8+ T-cell clones following secondary acute virus infection. Proc Natl Acad Sci U S A 111, 1485–1490 (2014).

49. Gil, A., Yassai, M.B., Naumov, Y.N. & Selin, L.K. Narrowing of human influenza A virus-specific T cell receptor alpha and beta repertoires with increasing age. J Virol 89, 4102–4116 (2015).

50. Selin, L.K., et al. Memory of mice and men: CD8+ T-cell cross-reactivity and heterologous immunity. Immunol Rev 211, 164–181 (2006).

51. Selin, L.K., et al. Heterologous immunity: immunopathology, autoimmunity and protection during viral infections. Autoimmunity 44, 328–347 (2011).

52. Benn, C.S., Netea, M.G., Selin, L.K. & Aaby, P. A small jab - a big effect: nonspecific immunomodulation by vaccines. Trends Immunol 34, 431–439 (2013).

53. Han, A., Glanville, J., Hansmann, L. & Davis, M.M. Linking T-cell receptor sequence to functional phenotype at the single-cell level. Nat Biotechnol 32, 684–692 (2014).

54. Bailey, T.L. & Elkan, C. Fitting a mixture model by expectation maximization to discover motifs in biopolymers. Proc Int Conf Intell Syst Mol Biol 2, 28–36 (1994).

55. Luzuriaga, K., et al. Early therapy of vertical human immunodeficiency virus type 1 (HIV-1) infection: control of viral replication and absence of persistent HIV-1-specific immune responses. J Virol 74, 6984–6991 (2000).

56. Krzywinski, M., et al. Circos: an information aesthetic for comparative genomics. Genome Res 19, 1639–1645 (2009).

57. Wang, G.C., Dash, P., McCullers, J.A., Doherty, P.C. & Thomas, P.G. T cell receptor alphabeta diversity inversely correlates with pathogen-specific antibody levels in human cytomegalovirus infection. Sci Transl Med 4, 128ra142 (2012).

58. Lefranc, M.P. IMGT, the international ImMunoGeneTics database. Nucleic Acids Res 31, 307–310 (2003).

59. Precopio, M.L., Sullivan, J.L., Willard, C., Somasundaran, M. & Luzuriaga, K. Differential kinetics and specificity of EBV-specific CD4+ and CD8+ T cells during primary infection. J Immunol 170, 2590–2598 (2003).

60. Garboczi, D.N., Hung, D.T. & Wiley, D.C. HLA-A2-peptide complexes: refolding and crystallization of molecules expressed in Escherichia coli and complexed with single antigenic peptides. Proceedings of the National Academy of Sciences of the United States of America 89, 3429–3433 (1992).

61. Powell, H.R., Battye, T.G.G., Kontogiannis, L., Johnson, O. & Leslie, A.G.W. Integrating macromolecular X-ray diffraction data with the graphical user interface iMosflm. Nature protocols 12, 1310–1325 (2017).

62. McCoy, A.J., et al. Phaser crystallographic software. Journal of applied crystallography 40, 658–674 (2007).

63. Winn, M.D., et al. Overview of the CCP4 suite and current developments. Acta crystallographica. Section D, Biological crystallography 67, 235–242 (2011).

64. Emsley, P., Lohkamp, B., Scott, W.G. & Cowtan, K. Features and development of Coot. Acta crystallographica. Section D, Biological crystallography 66, 486–501 (2010).

65. Adams, P.D., et al. PHENIX: a comprehensive Python-based system for macromolecular structure solution. Acta crystallographica. Section D, Biological crystallography 66, 213–221 (2010).

66. Karplus, P.A. & Diederichs, K. Linking crystallographic model and data quality. Science 336, 1030–1033 (2012).

